# mokapot: Fast and flexible semi-supervised learning for peptide detection

**DOI:** 10.1101/2020.12.01.407270

**Authors:** William E Fondrie, William S Noble

## Abstract

Proteomics studies rely on the accurate assignment of peptides to the acquired tandem mass spectra—a task where machine learning algorithms have proven invaluable. We describe mokapot, which provides a flexible semi-supervised learning algorithm that allows for highly customized analyses. We demonstrate some of the unique features of mokapot by improving the detection of RNA-cross-linked peptides from an analysis of RNA-binding proteins and increasing the consistency of peptide detection in a single-cell proteomics study.

## Main

Proteomics technologies seek to characterize the full complement of proteins in complex mixtures and have become invaluable for fields ranging from precision medicine to systems biology. Mass spectrometry is often the chosen method to detect and quantify the peptides derived from the enzymatic digest of complex protein mixtures, yielding insights into the abundance of the original proteins and their post-translational modifications. Critical to the success of mass spectrometry-based proteomics experiments is the accurate assignment of peptide sequences to the acquired mass spectra. The resulting peptide-spectrum matches (PSMs) are the foundation for inferring and quantifying the detected peptides and proteins [1].

The most common method to assign peptides to mass spectra is through a database search [2]. Proteomics search engines compare the theoretical fragments of peptide sequences from a sequence database against the acquired mass spectra, yielding one or more scores for each putative peptide-spectrum match (PSM). These scores quantify the quality of each PSM. Importantly, the incorporation of shuffled or reversed decoy peptide sequences into the sequence database allows for the accurate assignment of statistical confidence estimates to the selected set of PSMs [3]. Although many search engines perform well when PSMs are ranked by the search engine’s score function, often the sensitivity of peptide detection can be improved by integrating complementary scores and properties—“features”—that characterize a PSM.

Machine learning has been immensely successful at providing adaptable and unbiased methods to aggregate multiple features into a single score that greatly increases the sensitivity of peptide detection [4–6]. One such method, Percolator, introduced a method to learn models directly from the PSMs being analyzed. Percolator uses a set of confident PSMs as positive examples and decoy PSMs as negative examples to iteratively learn a support vector machine (SVM) that discriminates between them. The method is semi-supervised because the decoys have negative labels but the target labels must be inferred. Since its introduction, Percolator has been widely used and has demonstrably improved the ability to detect peptides with a wide variety search engines [7].

Despite its success, analyses conducted with Percolator are relatively rigid and may be suboptimal for certain types of experiments. For example, Percolator is limited to learning linear models, although it has been demonstrated that non-linear models can be beneficial [8]. Additionally, Percolator is intended to analyze each experiment independently; thus, to share a model across multiple experiments—such as when combining many experiments in a single-cell proteomics study—an external training dataset must be used to learn a static model[9].

Here we present mokapot, a fast and extensible Python implementation of Percolator’s semi-supervised algorithm that provides immense flexibility for highly customized analyses. We demonstrate the benefits of this flexibility in two vignettes: the detection of modified peptides from proteins cross-linked to RNA using an open modification search and the analysis of single-cell proteomics experiments.

Many proteomics experiments aim to characterize the post-translational modifications borne by proteins in a sample, and open modification searching [10]—where a peptide sequence is assigned to a mass spectrum while allowing for unspecified modifications—has become a popular method to accomplish this task. In this type of analysis, every PSM has an associated mass shift that indicates the difference between the observed precursor mass and the theoretical mass of the peptide. To demonstrate the utility of mokapot in this context, we analyzed a dataset where RNA-binding proteins were cross-linked to interacting RNA and subsequently analyzed by mass spectrometry [11]. After performing an open modification search with MSFragger [12], we were interested only in the modified peptides where we expected to find mass shifts corresponding to cross-linked RNA polymers. This analysis provided the opportunity to demonstrate two features of mokapot: grouped confidence estimates and the ability to use any type of machine learning classifier.

Grouped confidence estimates have proven to be useful when a subset of peptides are of interest or there is an expected shift in the score distribution between sets of PSMs in the experiment [13, 14]. In mokapot, PSMs can be easily assigned to groups, such that a model is learned from all of the PSMs, but confidence estimates are calculated separately within each group. For our analysis of the RNA-binding protein dataset, we were only interested in discovering modified peptides; hence, we designated PSMs with a mass shift greater than 50 ppm to be our group of interest.

The mass shift presents an interesting potential feature for models to learn from in mokapot. Although the mass shift can take any continuous value within the mass tolerance of our database search, we expect the correctly assigned PSMs to exhibit a discrete set of mass shifts corresponding to the masses of possible chemical moieties. Unfortunately, a linear model, such as the linear SVM employed by Percolator, cannot fully exploit the discrete nature of the mass shifts. However, we designed mokapot with a Scikit-learn interface [15] that makes it compatible with any type of machine learning classifier, allowing us to use non-linear models that can fully exploit the mass shift feature.

We analyzed the RNA-binding protein dataset using both the default linear SVM and a non-linear gradient boosting classifier implemented by XGBoost [16]. We found that the XGBoost classifier increased our sensitivity, enabling us to detect an additional 2,161 modified PSMs (15%, Figure 1a), 881 peptides (19%, Figure 1b), and 82 proteins (11%, Figure 1c) at a 1% false discovery rate (FDR). Furthermore, when we inspected the relative importance of each feature, we found that the XGBoost classifier relied more heavily on the mass shift feature than did the linear SVM (Supplementary Figure 1). We then investigated the mass shifts associated with PSMs gained or lost by the XGBoost classifier. We found a notable increase in the number of putative phosphorylated and carbamylated peptides detected (Figure 1c), consistent with the employed sample preparation procedure. More importantly, we also observed an increase in the number of PSMs with mass shifts corresponding to putative AU and ACU RNA cross-links.

**Figure 1:**
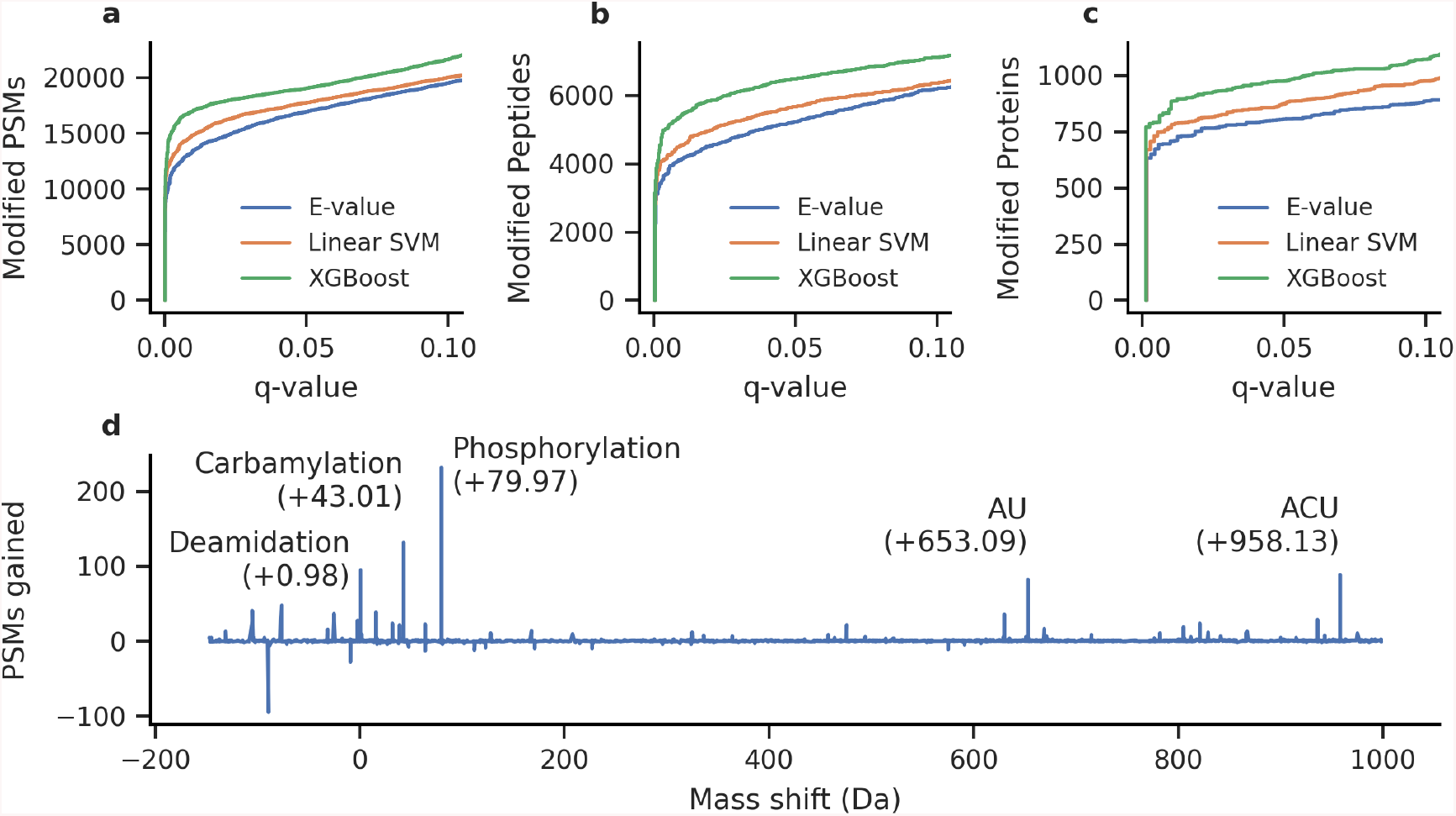
Mokapot improves the detection of RNA-cross-linked peptides from open modification search results. The non-linear XGBoost classifier resulted in the detection of more modified (a) PSMs, (b) peptides, and (c) proteins over a linear SVM (the default model in mokapot) or the MSFragger E-value. (d) The XGBoost classifier gained PSMs over the linear SVM with mass shifts that correspond to known modifications at 1% FDR.

Another common task for proteomics studies is the quantitative comparison of peptides and proteins detected from multiple experimental conditions. Although each experiment can be analyzed independently with mokapot, we previously observed increased variability and loss of power to detect PSMs and peptides when the individual experiments consist of relatively few total or confident PSMs [9]. One solution is to learn a static model from a large training dataset, then use the learned model to evaluate the small-scale experiments of interest. However, this static modeling approach relies on the availability of a training dataset that is independent of the experiments of interest, thereby restricting this approach to cases where a training set is available. Alternatively, mokapot offers a joint modeling approach: learn a joint model from the aggregate of all PSMs in a dataset, then assign confidence estimates within each experiment. We hypothesized that this joint modeling approach would offer a similar gain in power and consistency as observed with the static model, but without the need for an external training dataset.

We tested the joint modeling approach in mokapot by analyzing the single-cell proteomics dataset from Specht *et al.* [17] We sought to evaluate whether joint models improved the consistency of peptide detection across 65 experiments when compared against analyzing each experiment independently or using a static model. We found that the joint modeling approach consistently increased the numbers of confidently detected PSMs, peptides, and proteins at 1% FDR over analyzing each experiment independently (Figure 2A). Furthermore, the joint models increased the number of peptides and proteins detected across multiple experiments (Figure 2B and C), which is critical for reducing missing values in downstream quantitative analyses.

**Figure 2:**
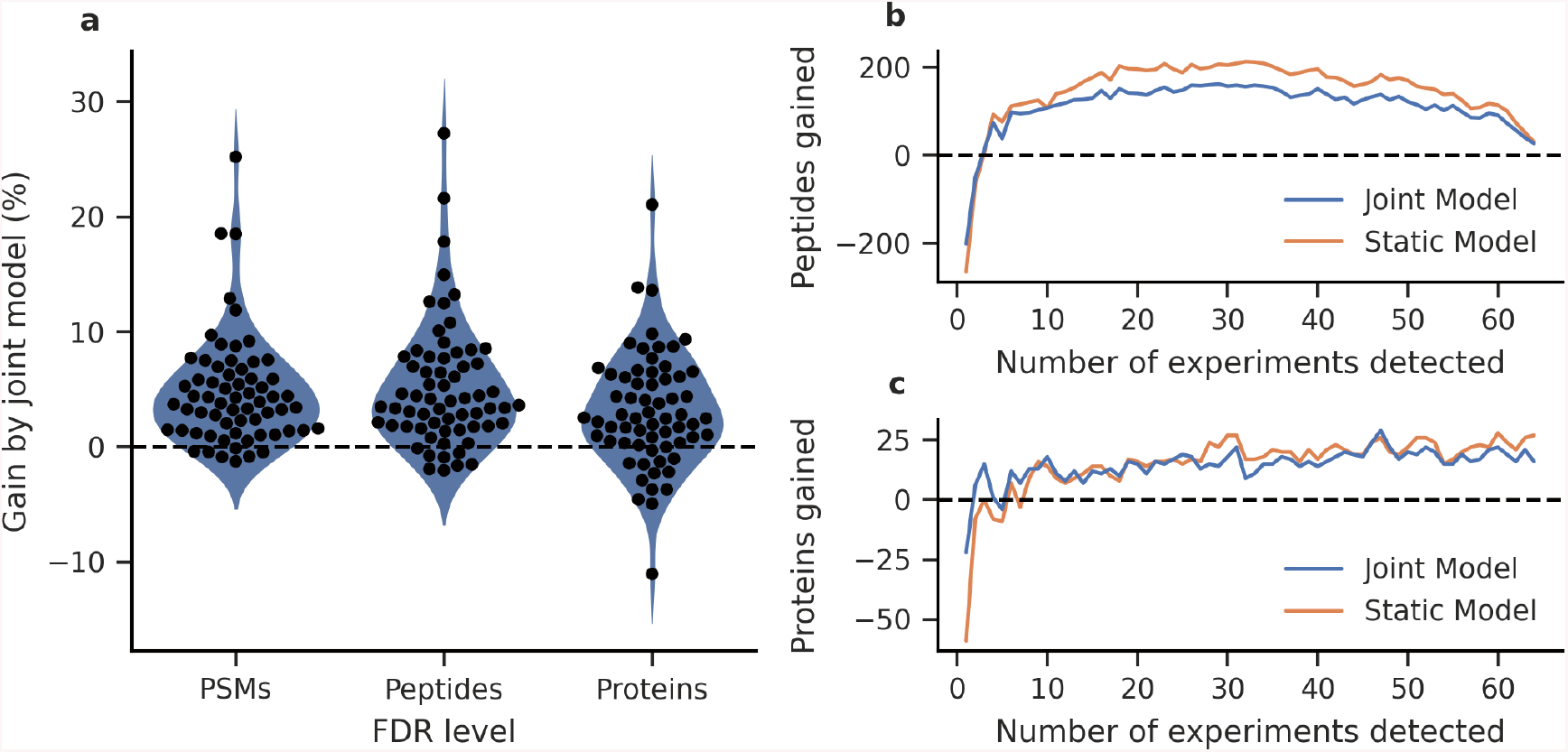
Joint models improve the power and consistency of peptide detection from single-cell proteomics experiments. (a) Joint models detect more PSMs, peptides, and proteins at 1% FDR than when experiments are analyzed individually. The detected (b) peptides and (c) proteins are more consistent across experiments using joint models in comparison to analyzing each experiment individually. In both cases, the joint models are comparable to using a static model, but without the requirement of a training dataset.

To be practically useful, mokapot must be fast. We profiled the runtime and memory usage of mokapot to assess its scalability. To accomplish this, we ran mokapot and Percolator on increasing numbers of PSMs sampled from the Kim *et al.* human proteome dataset [18]. These benchmarking experiments revealed that the runtime and memory usage of mokapot, like Percolator, scale approximately linearly with the number of PSMs when a linear SVM is used (Supplementary Figure 2). Additionally, we found that mokapot can reliably reproduce Percolator results when both are provided the same set of PSMs to analyze (Supplementary Figure 3).

Mokapot is an open-source Python package (https://github.com/wfondrie/mokapot) and can be readily extended to new types of proteomics data as they arise.

## Methods

### Benchmarking

We used the Kim *et al.* draft map of the human proteome dataset [18] to benchmark the performance of mokapot. This dataset consists of approximately 25 million mass spectra from LTQ-Orbitrap Elite and Velos mass spectrometers. The raw data files were downloaded from PRIDE [19] (PXD000561) and converted to ms2 format using msconvert [20] with peak-picking and deisotoping filters (“peakPicking vendor msLevel=2” and “MS2Deisotope Poisson”).

We searched the converted files against the canonical UniProt [21] human proteome using the Tide search engine [22] with the combined p-value score function [23]. Trypsin without proline suppression was used, allowing for two missed cleavages. Variable methionine oxidation and protein N-terminal acetylation modifications were allowed. Additionally, carbamidomethylation of cysteine was specified as a static modification. We selected the precursor *m/z* window using Param-Medic [24] and set a fragment ion tolerance of 0.02 Da. The protein database was processed using Tide to generate a shuffled decoy peptide sequence for each peptide sequence in the target database, preserving both termini.

We compared the scores and confidence estimates of mokapot to Percolator v3.05 when used to analyze the same random sample of 1 million PSMs from the full Kim *et al.* dataset. Additionally, we benchmarked the speed and memory usage of mokapot and Percolator on various sets of PSMs sampled from the full dataset across logarithmically spaced intervals. We used the GNU time application to record the total run time and the maximum memory used (the maximum resident set size) for both applications run as command line tools. We repeated this process three times for each set of PSMs and reported the minimum. These analyses were conducted on a cluster node running CentOS7 and equipped with 12 CPU cores (Intel Xeon E5-2680v3) and 96 Gb of memory.

### Cross-linked RNA-binding proteins

Kramer *et al.* [11] provide multiple datasets created from RNA-binding proteins cross-linked to interacting RNA molecules using ultraviolet radiation. We chose to reanalyze the yeast experiments from this study as an example of how mokapot can enhance the detection of modified peptides. The raw data files were downloaded from PRIDE Archive (PXD000513) and converted to mzML format using ThermoRawFileParser [25], with vendor peak-picking enabled.

We searched the converted files against the canonical UniProt yeast (*Saccharomyces cerevisiae* [strain ATCC 204508 / S288c]) reference proteome concatenated with decoy sequences (6,721 target protein sequences, downloaded November 4, 2020) using MSFragger v3.1.1 [12]. Decoy sequences were generated with mokapot by shuffling non-terminal amino acids within each tryptic peptide sequence. We performed an open modification search using a precursor mass window from −150 to 1000 Da, with mass calibration and modification localization enabled [26]. We used trypsin without proline suppression as the enzyme specificity and allowed for two missed cleavages. Carbamidomethylation of cysteine was not allowed. Methionine oxidation and protein N-terminal acetylation were specified as variable modifications. We allowed the top five matches per spectrum to be reported, so that mokapot would have the opportunity to re-rank them, resulting in 382,148 total PSMs.

We made minor modifications to the Percolator input files that were generated by MSFragger. First, the original *ExpMass* column was renamed to *CalcMass*, because it contained the theoretical peptide mass. The neutral precursor mass was then added as a new *ExpMass* column. These changes were necessary to ensure that target-decoy competition resulted in a single match per spectrum after mokapot analysis. The observed mass shifts were binned into 0.01 Da increments and appended to the sequence string in the *Peptide* column to define modified peptides for confidence estimation. Additionally, we removed the *delta_hyperscore* feature out of an abundance of caution for excluding potential bias, particularly for our non-linear models. Finally, we added a *group* column to indicate whether a peptide was modified—which we defined as having a mass shift greater than 50 ppm—to serve as the groups for grouped confidence estimation within mokapot. The features used for mokapot are detailed in Supplementary Table 1.

Ranking PSMs by the MSFragger E-value was used as a baseline to compare against mokapot’s performance using a linear SVM or a non-linear XGBoost classifier [16]. For the XGBoost classifier, mokapot allowed us to perform a hyperparameter grid search using the nested cross-validation strategy proposed by Granholm *et al* [27] over the following parameters: *scale_pos_weight* (1, 10, 100), *max_depth* (1, 3, 6), *min_child_weight* (1, 10, 100), and *gamma* (0, 1, 10). The feature importance for both models were estimated by randomly permuting each feature and observing the average effect on the score for all modified PSMs, repeated five times [28]. These values were then normalized such that the sum of all features was one, providing a relative scale to compare the two models.

### Single-cell proteomics

We chose to analyze the Specht *et al.* single-cell proteomics dataset [17] to demonstrate the benefits that joint modeling provides. This dataset consists of two sets of experiments: a set of 76 mass spectrometry acquisitions for quality control and a set of 69 mass spectrometry acquisitions assessing macrophage differentiation. All of the acquisitions were performed on a Q-Exactive mass spectrometer, where a single acquisition is a multiplexed experiment analyzing multiple single cells using tandem mass tag (TMT) 10-plex reagents. The raw data files were downloaded from MassIVE [29] (MSV000083945) and converted to mzML format using ThermoRawFileParser, with vendor peak-picking enabled.

We searched the converted files against the canonical UniProt/Swiss-Prot human proteome (20,416 protein sequences, downloaded September 6, 2019) using Tide with the combined p-value score function [23]. Trypsin without proline suppression was used, allowing for two missed cleavages. The protein database was processed using Tide to generate a shuffled decoy peptide sequence for each peptide sequence in the target database, preserving both termini. We selected the precursor *m/z* window using Param-Medic [24] and set a fragment ion tolerance of 0.02 Da. We included the TMT 10-plex modification of lysine and the peptide N-terminus as a static modification, but carbamidomethylation of cysteine was not included. Additionally, we considered the oxidation of methionine, protein N-terminal acetylation, and deamidation of asparagine as variable modifications. Four of the macrophage differentiation experiments resulted in no PSMs at a 1% FDR threshold and were excluded from further analysis.

We analyzed the macrophage differentiation experiments with mokapot using three approaches: treating each experiment independently, learning a joint model from all of the macrophage differentiation experiments, or learning a static model from the quality control experiments and applying it to the macrophage differentiation experiments.

### The mokapot algorithm

The algorithm for the primary mokapot workflow begins by dividing the provided PSMs into cross-validation splits as proposed by Granholm *et al.* [27] and implemented in Percolator. Each split is comprised of a disjoint training set and test set, where the training set is used to train the model. The final score for each PSM is derived from the score assigned by the learned model to that PSM only when it is a member of the test set; thus, no PSMs are scored by the same model that they were used to train.

Model training occurs in accordance with the Percolator algorithm. First, positive examples are defined as PSMs in the training set that meet a prescribed FDR threshold using the best feature—that is, the feature that yields the largest number of discoveries at the FDR threshold provided when used to rank the PSMs. Likewise, negative examples are defined as decoy PSMs in the training set. Optionally, a hyperparameter optimization method can be employed within each split before proceeding to train the model. By default, mokapot uses a linear SVM as its model and employs a hyperparameter grid search to determine the cost of positive and negative misclassifications, emulating the approach used by Percolator. However, any model and hyperparameter optimization approach compliant with the Scikit-learn interface can be used in mokapot.

Model training then proceeds in an iterative manner. In each iteration, the model learns to separate the positive and negative examples it is provided. Once complete, the model is used re-score all of the PSMs in the training set, and a new set of positive examples is defined by assessing which PSMs meet the prescribed FDR threshold when ranked by the new score. This process is repeated, aggregating more PSMs into the positive examples with each iteration for typically ten iterations or until the selected set of positives stops changing.

After a model is trained with each cross-validation split, the learned models are used to score their respective test sets. When the model does not already output a calibrated score—such as a probability—mokapot calibrates the score between cross-validation splits, again according to Granholm *et al.* [27]. Finally, confidence estimation is performed with PSMs ranked by the scores provided by the new models. When a joint model is used, confidence estimates are assigned independently for each experiment. Furthermore, if groups are defined, then confidences estimates are assigned independently within each group as well [13, 14].

### Confidence estimation

Mokapot provides two forms of confidence estimates: q-values and posterior error probabilities (PEPs). The q-value is defined as the minimal false discovery rate (FDR) threshold at which a discovery would be accepted. In mokapot, the FDR for a collection of PSMs and peptides is estimated using target-decoy competition. Formally, we denote the number of target discoveries as *m_t_* and the number of decoy discoveries as *m_d_*. Likewise, we denote scores of target discoveries as 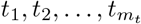 and the scores of decoy discoveries as 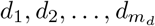. For each spectrum, we retain only the best scoring peptide, with ties broken randomly so as not to impose a bias toward target or decoy PSMs. For peptide-level confidence estimation, we retain only the best scoring PSM per peptide. The estimated FDR at a threshold *τ* is computed as the number of decoy discoveries that meet the threshold—which provide an estimate for the number of false positives accepted at this threshold—divided by the number of target discoveries that meet the threshold. We add one to the number of decoy discoveries, so as to prevent liberal FDR estimates for smaller collections of discoveries:

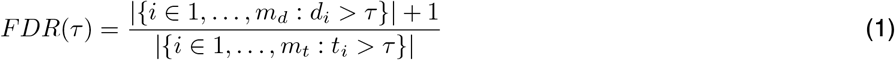

Thereafter, the q-value is defined as:

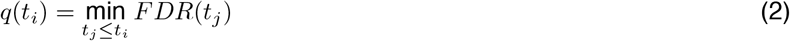

Protein-level q-values are estimated using the picked-protein approach [30]. Briefly, protein sequences are assigned to proteins groups if the enzymatic peptides form a formal subset of those generated by another protein. The best PSM for each protein group is then retained and target proteins are competed against their decoy counterparts, eliminating the lower scoring of the pair. The protein groups are then assigned q-values using equations 1–2.

The PEP is the probability that the observed discovery is incorrect. In mokapot, we use the qvality method [31] as implemented in triqler [32] to estimate the PEP for each PSM, peptide, and protein group after the competition procedures described above.

## Data and code availability

Mokapot is an open source project and is publicly available on GitHub (https://github.com/wfondrie/mokapot). All of the datasets we used throughout this study are available through their respective ProteomeXchange [33] partner repositories. All code used for these analyses and to generate the figures presented in this study is available on GitHub (https://github.com/Noble-Lab/mokapot-analyses).

## Acknowledgments

We are grateful to Lukas Käll and Matthew The for helpful discussions about the design of mokapot. The research reported in this publication was supported by the National Institutes of Health awards T32HG000035, P41GM103533, and R01GM121818.

## References

[1] Hunt, D. F., Yates, III, J. R., Shabanowitz, J., Winston, S., et al. “Protein sequencing by tandem mass spectrometry.” In: Proceedings of the National Academy of Sciences of the United States of America 83 (1986), pp. 6233–6237.

[2] Eng, J. K., McCormack, A. L., and Yates, III, J. R. “An approach to correlate tandem mass spectral data of peptides with amino acid sequences in a protein database.” In: Journal of the American Society for Mass Spectrometry 5 (1994), pp. 976–989.

[3] Elias, J. E. and Gygi, S. P. “Target-decoy search strategy for increased confidence in large-scale protein identifications by mass spectrometry.” In: Nature Methods 4.3 (2007), pp. 207–214.

[4] Keller, A., Nesvizhskii, A. I., Kolker, E., and Aebersold, R. “Empirical statistical model to estimate the accuracy of peptide identification made by MS/MS and database search.” In: Analytical Chemistry 74 (2002), pp. 5383–5392.

[5] Anderson, D. C., Li, W., Payan, D. G., and Noble, W. S. “A new algorithm for the evaluation of shotgun peptide sequencing in proteomics: support vector machine classification of peptide MS/MS spectra and Sequest scores.” In: Journal of Proteome Research 2.2 (2003), pp. 137–146.

[6] Käll, L., Canterbury, J., Weston, J., Noble, W. S., et al. “A semi-supervised machine learning technique for peptide identification from shotgun proteomics datasets.” In: Nature Methods 4 (2007), pp. 923–25.

[7] Tu, C., Sheng, Q., Li, J., Ma, D., et al. “Optimization of search engines and postprocessing approaches to maximize peptide and protein identification for high-resolution mass data.” In: Journal of Proteome Research 14.11 (2015), pp. 4662–4673.

[8] Spivak, M., Weston, J., Bottou, L., Käll, L., et al. “Improvements to the Percolator algorithm for peptide identification from shotgun proteomics data sets.” In: Journal of Proteome Research 8.7 (2009), pp. 3737–3745.

[9] Fondrie, W. E. and Noble, W. S. “Machine Learning Strategy That Leverages Large Data Sets to Boost Statistical Power in Small-Scale Experiments.” eng. In: Journal of Proteome Research 19.3 (Mar. 2020), pp. 1267–1274.

[10] Chick, J. M., Kolippakkam, D., Nusinow, D. P., Zhai, B., et al. “A mass-tolerant database search identifies a large proportion of unassigned spectra in shotgun proteomics as modified peptides.” In: Nature Biotechnology (2015). Epub ahead of print.

[11] Kramer, K., Sachsenberg, T., Beckmann, B. M., Qamar, S., et al. “Photo-Cross-Linking and High-Resolution Mass Spectrometry for Assignment of RNA-Binding Sites in RNA-Binding Proteins.” en. In: Nature Methods 11.10 (Oct. 2014), pp. 1064–1070.

[12] Kong, A. T., Leprevost, F. V., Avtonomov, D. M., Mellacheruvu, D., et al. “MSFragger: ultrafast and comprehensive peptide identification in mass spectrometry-based proteomics.” In: Nature Methods 14.5 (2017), pp. 513–520.

[13] Efron, B. “Microarrays, Empirical Bayes and the Two-Groups Model.” In: Statistical Science 23.1 (2008), pp. 1–22.

[14] Yi, X., Gong, F., and Fu, Y. “Transfer posterior error probability estimation for peptide identification.” In: BMC Bioinformatics 21 (May 2020).

[15] Pedregosa, F., Varoquaux, G., Gramfort, A., Michel, V., et al. “Scikit-learn: Machine Learning in Python.” In: Journal of Machine Learning Research 12 (2011), pp. 2825–2830.

[16] Chen, T. and Guestrin, C. “XGBoost: A Scalable Tree Boosting System.” In: Proceedings of the 22nd ACM SIGKDD International Conference on Knowledge Discovery and Data Mining. KDD ’16. San Francisco, California, USA: ACM, 2016, pp. 785–794.

[17] Specht, H., Emmott, E., Koller, T., and Slavov, N. “High-throughput single-cell proteomics quantifies the emergence of macrophage heterogeneity.” In: bioRxiv (2019).

[18] Kim, M., Pinto, S. M., Getnet, D., Nirujogi, R. S., et al. “A draft map of the human proteome.” In: Nature 509.7502 (2014), pp. 575–581.

[19] Vizcaíno, J. A., Csordas, A., del-Toro, N., Dianes, J. A., et al. “2016 update of the PRIDE database and its related tools.” In: Nucleic Acids Research 44 (D1 2016), pp. D447–D456.

[20] Kessner, D., Chambers, M., Burke, R., Agnus, D., et al. “ProteoWizard: open source software for rapid proteomics tools development.” In: Bioinformatics 24.21 (2008), pp. 2534–2536.

[21] The UniProt Consortium. “UniProt: a worldwide hub for protein knowledge.” In: Nucleic Acids Research (2019), pp. D506–D515.

[22] Diament, B. and Noble, W. S. “Faster SEQUEST searching for peptide identification from tandem mass spectra.” In: Journal of Proteome Research 10.9 (2011), pp. 3871–3879.

[23] Lin, A., Howbert, J. J., and Noble, W. S. “Combining High-Resolution and Exact Calibration To Boost Statistical Power: A Well-Calibrated Score Function for High-Resolution MS2 Data.” In: Journal of Proteome Research 17 (11 2018), pp. 3644–3656.

[24] May, D. H., Tamura, K., and Noble, W. S. “Param-Medic: A tool for improving MS/MS database search yield by optimizing parameter settings.” In: Journal of Proteome Research 16.4 (2017), pp. 1817–1824.

[25] Hulstaert, N., Sachsenberg, T., Walzer, M., Barsnes, H., et al. “ThermoRawFileParser: modular, scalable and cross-platform RAW file conversion.” In: Journal of Proteome Research 19.1 (2020), pp. 537–542.

[26] Yu, F., Teo, G. C., Kong, A. T., Haynes, S. E., et al. “Identification of Modified Peptides Using Localization-Aware Open Search.” en. In: Nature Communications 11.1 (Aug. 2020), p. 4065.

[27] Granholm, V., Noble, W. S., and Käll, L. “A cross-validation scheme for machine learning algorithms in shotgun proteomics.” In: BMC Bioinformatics 13.Suppl 16 (2012), S3.

[28] Breiman, L. “Random forests.” In: Machine Learning 45.1 (2001), pp. 5–32.

[29] Wang, M., Wang, J., Carver, J., Pullman, B. S., et al. “Assembling the Community-Scale Discoverable Human Proteome.” In: Cell Systems 7 (4 2018), 412–421.e5.

[30] Savitski, M. M., Wilhelm, M., Hahne, H., Kuster, B., et al. “A scalable approach for protein false discovery rate estimation in large proteomic data sets.” In: Molecular & Cellular Proteomics 14.9 (2015), pp. 2394–2404.

[31] Käll, L., Storey, J. D., and Noble, W. S. “qvality: Nonparametric estimation of *q* values and posterior error probabilities.” In: Bioinformatics 25.7 (2009), pp. 964–966.

[32] The, M. and Käll, L. “Integrated Identification and Quantification Error Probabilities for Shotgun Proteomics.” en. In: Molecular & Cellular Proteomics 18.3 (Mar. 2019), pp. 561–570.

[33] Vizcaíno, J. A., Deutsch, E. W., Wang, R., Csordas, A., et al. “ProteomeXchange provides globally coordinated proteomics data submission and dissemination.” In: Nature Biotechnology 32 (2014), pp. 223–226.

