## Supplemental Information for "mokapot: Fast and flexible semi-supervised learning for peptide detection"

| # | Feature | Description |
|---|---|---|
| 1 | rank | The rank of the PSM for each spectrum by MSFragger |
| 2 | abs_ppm | The absolute value of the mass shift in parts-per-million |
| 3 | abs_mass_diff | The absolute value of the mass shift in Daltons |
| 4 | log10_evalue | The $log_{10}$ of the MSFragger E-Value |
| 5 | hyperscore | The MSFragger hyperscore |
| 6 | matched_ion_num | The number of matched theoretical ions |
| 7 | matched_ion_fraction | The fraction of matched theoretical ions |
| 8 | peptide_length | The length of the peptide |
| 9 | ntt | The number of enzymatic termini |
| 10 | nmc | The number of missed cleavages |
| 11–17 | charge_[1–7] | Seven Boolean features indicating charge state |

**Supplementary Table 1:** The MSFragger features used by mokapot

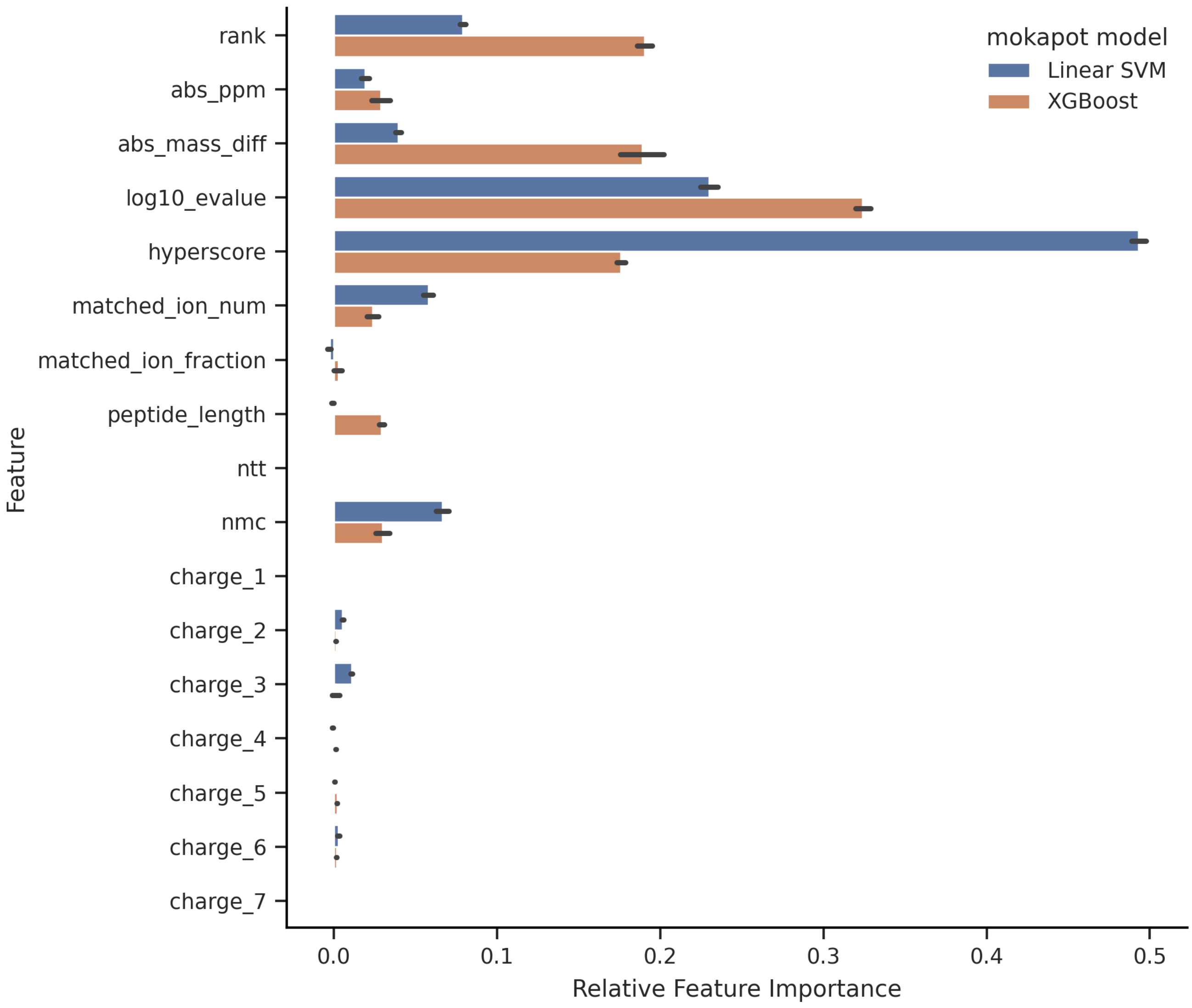

**Supplementary Figure 1:** The relative feature importance of a linear SVM and non-linear XGBoost classifier toward the PSM scores in mokapot. In comparison to the linear SVM, the XGBoost classifier relied more heavily on the mass shift (abs_mass_diff), rank, peptide length (peptide_length), and the MSFragger E-value (log10_evalue), while relying less on the MSFragger hyperscore.

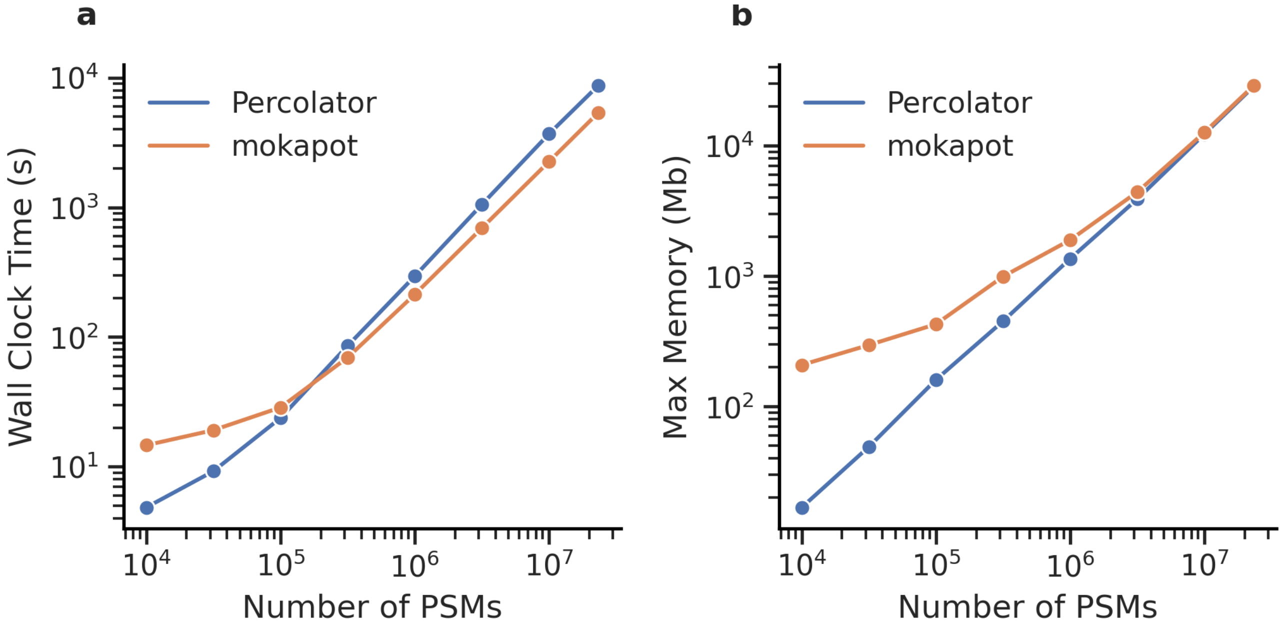

**Supplementary Figure 2:** Mokapot is fast and memory efficient. (a) The run time of mokapot scales favorably when compared to Percolator on large datasets. (b) Mokapot has a significant memory overhead for small datasets, but scales similarly to Percolator for large datsets.

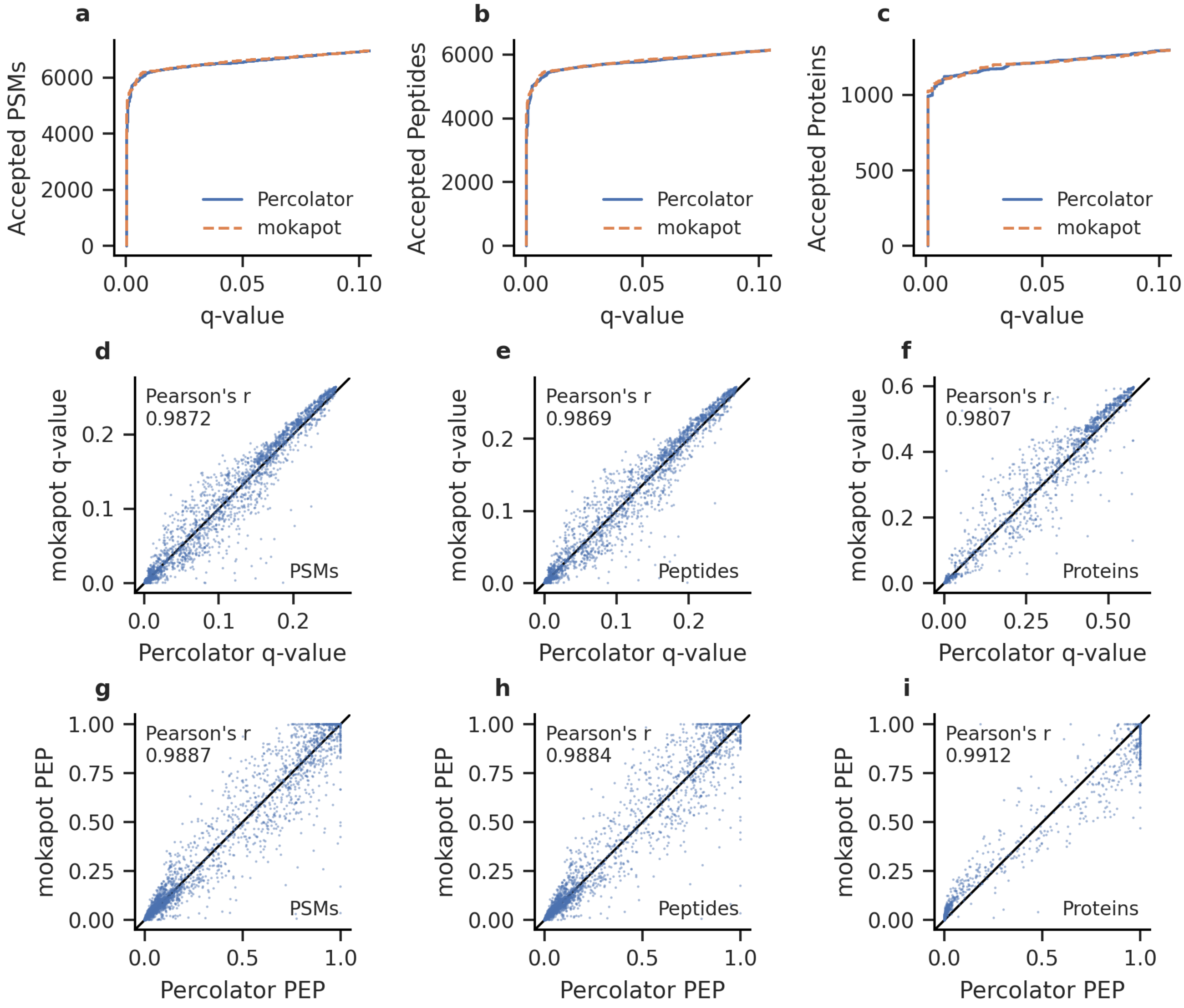

**Supplementary Figure 3:** Mokapot can reproduce Percolator analyses. When mokapot and Percolator were used to analyze a single-cell proteomics experiment, similar numbers of (a) PSMs, (b) peptides, and (c) proteins are accepted at low FDR thresholds. (d–f) The q-values reported by mokapot are highly correlated with those reported by Percolator. (g–i) Likewise, the posterior error probabilities reported by mokapot and Percolator are also highly correlated.
